# How do Large Language Models understand Genes and Cells

**DOI:** 10.1101/2024.03.23.586383

**Authors:** Chen Fang, Yidong Wang, Yunze Song, Qingqing Long, Wang Lu, Linghui Chen, Pengfei Wang, Guihai Feng, Yuanchun Zhou, Xin Li

**Affiliations:** State Key Laboratory of Stem Cell and Reproductive Biology, Institute of Zoology, Chinese Academy of Sciences and University of the Chinese Academy of Sciences, China; Peking University, China; University of Liverpool, UK; Computer Network Information Center, Chinese Academy of Sciences, China; Tsinghua University, China; iFlytek Co Ltd, China; State Key Laboratory of Stem Cell and Reproductive Biology, Institute of Zoology, Chinese Academy of Sciences, China

**Author notes:** Both authors contributed equally to this research (Chen Fang, Yidong Wang).

**Keywords:** large language models, cell biology, gene gene interaction, cell annotation

## Abstract

Researching genes and their interactions is crucial for deciphering the fundamental laws of biological activity, advancing disease treatment, drug discovery and so on. Large language Models (LLMs), with their profound text comprehension and generation capabilities, have made significant strides across various natural science fields. However, their application in cell biology remains notably scarce. To alleviate this issue, in this paper, we selects seven mainstream LLMs and evaluates their performance across a range of problem scenarios. Our findings indicate that LLMs possess a certain level of understanding of genes and cells, and hold potential for solving real-world problems. Moreover, we have improved the current method of textual representation of cells, enhancing the LLMs’ ability to tackle cell annotation tasks. We encourage cell biology researchers to leverage LLMs for problem-solving while also being mindful of some challenges associated with their use. We release our code and data at https://github.com/epang-ucas/Evaluate_LLMs_to_Genes.

**CCS Concepts:** Applied computing → Recognition of genes and regulatory elements; Bioinformatics; Computational genomics; Computational transcriptomics.

**ACM Reference Format:** Chen Fang, Yidong Wang, Yunze Song, Qingqing Long, Wang Lu, Linghui Chen, Pengfei Wang, Guihai Feng, Yuanchun Zhou, and Xin Li. 2024. How do Large Language Models understand Genes and Cells. 1, 1 (March 2024), 14 pages. https://doi.org/10.1145/nnnnnnn.nnnnnnn

## 1 INTRODUCTION

Genes are the fundamental units within organisms that control the transmission and expression of hereditary traits. Investigating the functions of genes and their interrelationships is crucial for unveiling the genetic and developmental mechanisms of organisms. This understanding is instrumental in identifying the root causes of genetic diseases and in the development of novel therapeutic approaches. Cells can be seen as collections of genes. The selective expression of these genes dictates the direction of cell differentiation, thereby endowing cells with distinct morphologies and functions. The advancement of single-cell sequencing technologies[13, 15] has enabled us to observe gene expression from the perspective of individual cells, accumulating vast amounts of transcriptomic data. This, in turn, facilitates the application of deep learning methods in scenarios related to genes and cells[19, 26, 30]. However, such methods heavily rely on the quality of transcriptional sequencing data, the collection and processing of which are notably time-consuming and labor-intensive.

The success of Large Language Models (LLMs) in the field of Natural Language Processing(NLP) suggests an alternative method for problem-solving. LLMs gain a general understanding of natural language through pre-training on vast corpora and then undergo fine-tuning for specific problem scenarios, thus aiding in solving practical issues. LLMs can be applied to various scientific research fields with just appropriate textual input (prompt), without the need for additional complex modeling. Nonetheless, the application of LLMs in addressing gene-related issues has been sparse: GenePT[5] utilizes LLMs to generate embedding vectors for genes and cells while Cell2Sentence[16] has fine-tuned GPT-2 for cell annotation tasks. To alleviate this issue, this paper focuses on exploring the performance of LLMs across a spectrum of gene-related problems and evaluates the effectiveness of several mainstream LLMs in these contexts. To comprehensively assess the understanding of LLMs on gene and cell-related issues, we examine their capabilities in three aspects: identification of genes, prediction of gene interrelations, and cell-level annotation tasks. In addition, both GenePT and Cell2Sentence independently employ a method where the names of the top 100 highly expressed genes are concatenated to form a textual representation of a cell, referred to as a “cell sentence”. However, we argue that such representations lack the textual structure characteristic of natural language. To make the cell sentence more comprehensible to LLMs, we have appended a succinct functional description to each gene name, a method we have dubbed “cell sentence plus”. Empirical evidence supports that this approach indeed enhances the performance of LLMs.

The contributions of this paper can be summarized as follows:

- We provide guidance for fine-tuning LLMs to address gene-related issues.
- We evaluate the performance of several mainstream LLMs in gene-related application scenarios.
- For cell annotation tasks, we have improved the mainstream textual representation of cells by adding textual descriptions after each gene name, which enhances the effectiveness of LLMs.
- Our training code, details, and data are open-sourced at https://github.com/epang-ucas/Evaluate_LLMs_to_Genes, which can facilitate future research on this topic on using LLMs for gene-related problems.

## 2 RELATED WORK

Recently, research in LLMs has experienced an explosive growth, pushing forward not only traditional tasks in NLP such as text classification, text generation, conversational bots and so on, but also accelerating the pace of innovation in research methodologies across the natural sciences. In this wave, several notable LLM projects have stood out. ChatGPT[22], developed by OpenAI, is a significant LLM based on the GPT[23] architecture, recognized for its conversation abilities akin to human interactions and its extensive knowledge comprehension. Meta’s LLaMA[34] aims to enhance training efficiency and lower computational expenses, highlighting its openness and accessibility. BLOOM[27], from the BigScience research workshop, is an open-source, multilingual LLM emphasizing its linguistic versatility and commitment to open scientific practices. THUDM’s ChatGLM[46] targets the Chinese market, focusing on generation capabilities within specific linguistic contexts. Anthropic’s Mistral[14] prioritizes interpretability, seeking to improve transparency and user comprehension of model behaviors. These examples represent leading efforts in the realm of universal LLM research, hence their selection for analysis in this paper. Furthermore, in the biosciences, BioGPT[20] is dedicated to generative pre-training in the biomedical field, aiming to process and generate text related to biomedicine, marking it as another focus of our evaluation.

The application of LLMs in solving gene-related problems in cell biology poses an ongoing challenge. GPTCelltype[11] employs GPT-4 to identify cell types based on marker gene information. Cell2Sentence[16], through fine-tuning GPT-2, annotates cells by generating a textual representation for each one, based on the names of the top 100 genes ranked by their transcriptional expression values in the cell. GenePT[5] uses a similar approach for textualizing cells and employs GPT-3.5 to generate embedding vectors, which are then combined with other supervised learning models for various downstream tasks. The scarcity of efforts in fine-tuning LLMs for gene-related issues is apparent, with only Cell2Sentence being a notable attempt at cell annotation. This paper explores the potential of LLMs in cell biology by fine-tuning seven LLMs across nine datasets, representing a diverse range of problem scenarios.

Beyond directly deploying LLMs to tackle gene-related issues, the success of LLMs has also sparked a trend in the field of cellular biology to construct specialized pre-trained foundation models[6, 9, 32, 42]. The underlying idea of these models is to draw an analogy between the concept of cells in biology to sentences in the field of NLP, and similarly, liken the concept of genes within cells to the words that make up sentences. This approach enables the application of transformer models to mine transcriptional expression data and to be utilized in a variety of downstream tasks such as cell annotation, inference of gene regulatory networks, and prediction of gene perturbations. In recent years, advances in high-throughput sequencing technologies along with efforts[1, 25, 41] in standardizing data collection and management have made it feasible to apply this “pre-training & fine-tuning” strategy on transcriptomics. These foundation models have indeed achieved the best results so far in specific problem scenarios. However, their demands for massive amounts of data and computational resources present significant challenges.

## 3 METHODS

In this paper, we select 7 generative LLMs and evaluate them across 9 datasets representing diverse problem scenarios. All evaluation tasks can be conceptualized to understand genes from three perspectives: individual genes, gene pairs, and cells. We employed the LORA[12] method uniformly across all LLMs for fine-tuning, with the implementation based on Llama-factory[10]. All fine-tuning experiments were performed on a single A800 GPU with a learning rate set to 0.00005. Figure 1 illustrates our process for fine-tuning LLMs to address gene-related issues comprehensively. Initially, we represented genes or cells in text form, followed by the application of appropriate prompts for fine-tuning the LLMs. This section will first describe our method for textual representation of genes and cells, then introduce the downstream tasks selected for evaluating the LLMs.

**Fig. 1.**
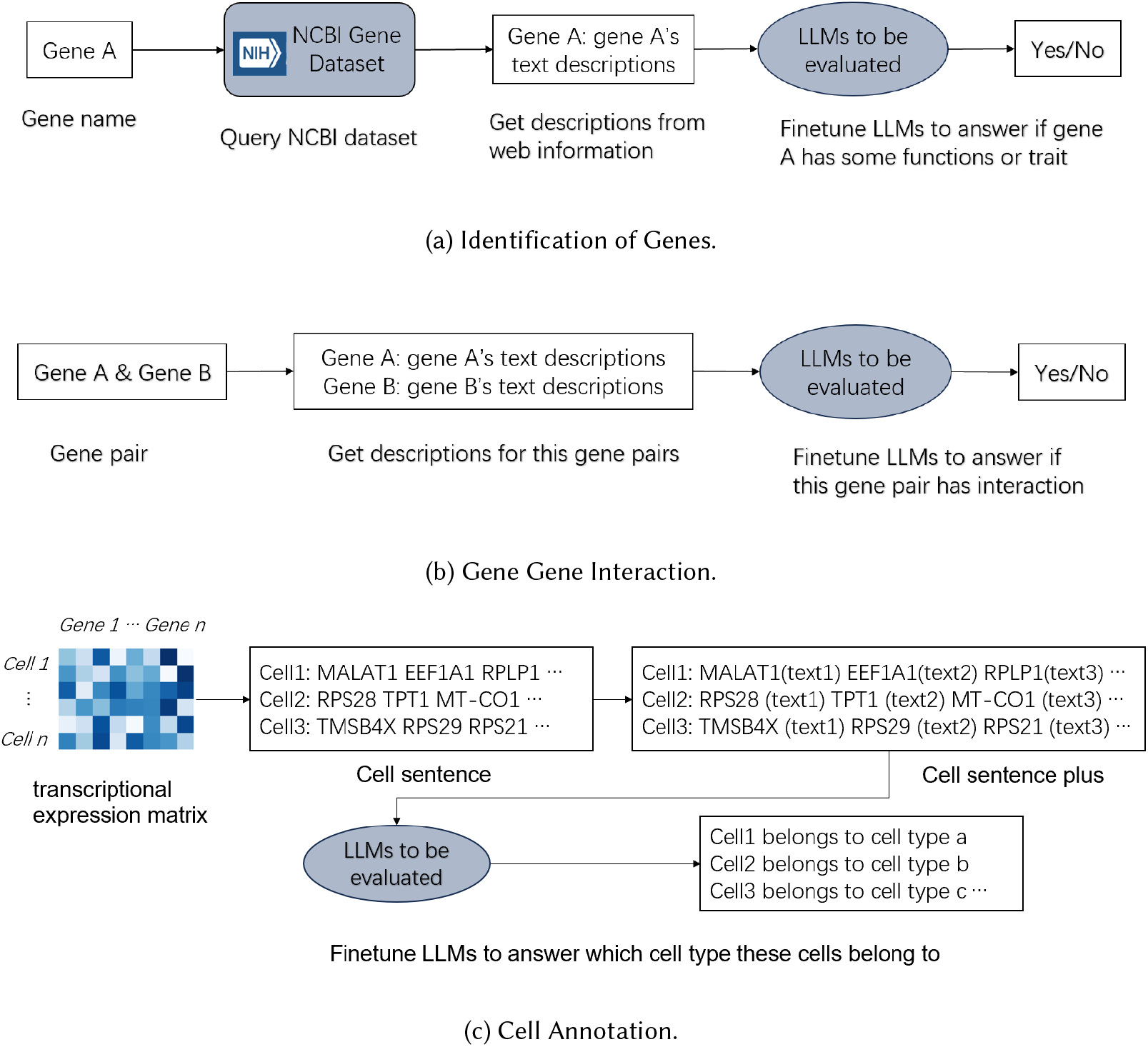
Overview of fine-tuning LLMs to solve gene-related problems. (a) For tasks with respect to the identification of genes, we first retrieve gene textual records from the NCBI database, then fine-tune LLMs to predict whether a gene has a certain function or trait. (b) For predicting the interaction between gene pairs, we input two genes and their respective NCBI descriptions, and fine-tune LLMs to determine if there is an interaction between them. (c) For cell annotation tasks, based on the expression values of genes within a cell, we select the top 100 genes to form a cell sentence. Then, by adding concise text descriptions after each gene name to create a cell sentence plus, we fine-tune LLMs to identify the cell type.

### 3.1 Textual Representation of Genes and Cells

To ensure LLMs fully comprehend the target genes, textual descriptions about these genes are necessary. The NCBI is a crucial resource offering an extensive array of databases and analytical tools in the biomedical and biotechnological fields. Its gene database[3] contains detailed records of the majority of genes, including their names, symbols, locations, gene products (such as proteins), and various attributes, such as gene functions, phenotypes, and interactions, among others. We utilized the complete records from NCBI as the textual description for each gene. For genes not included in the NCBI Gene Database, we generated a pseudo-description by consulting GPT-3.5 as an alternative.

Textual descriptions of cells are more complex. The transcriptional expression values of genes represent the absolute or relative quantity of transcriptional products under specific conditions. Typically, the transcriptional expression values of all genes within a cell are sufficient to reflect the cell’s identity and state. The mainstream approach for textualizing cells involves sorting all genes within a cell by their transcriptional expression values, retaining the top 100 genes, and sequentially concatenating their names. This method, known as “cell sentence” within Cell2Sentence[16], results in a sentence composed of 100 gene names without a natural linguistic structure, significantly diverging from the corpus LLMs encountered during their pre-training phase. To make the “cell sentence” more sentence-like, we introduced the “cell sentence plus”, where each gene name is followed by a brief description of that gene, facilitating the LLMs’ understanding of the input’s meaning, as illustrated in Figure 1c. Additionally, considering the lengthy descriptions of genes in NCBI, we employed GPT-3.5’s API service to refine all gene descriptions and limit their length to within 10 tokens, resulting in cell sentence plus lengths of approximately 2000 tokens, well within the input capacity of most LLMs (which typically have a maximum token limit of 4096).

### 3.2 Downstream Tasks Related to Genes and Cells

#### 3.2.1 Identification of Genes

The first category of tasks aims to explore the LLMs’ understanding of individual genes, examining whether they can extract and even infer specific knowledge about a gene based on records from the NCBI database. Following the setup of downstream tasks in GeneFormer[32], we evaluate the performance of LLMs across six datasets encompassing various aspects, including dosage sensitivity, chromatin dynamics, and network dynamics. To simplify these issues, all tasks have been formalized into binary classification tasks. We briefly described the task objectives to the LLMs, provided them with the gene under investigation along with its corresponding NCBI record, and queried whether the gene possesses a certain function or characteristic. An exemplary prompt template is illustrated in Figure 2. It is noteworthy that due to the high costs associated with biological experimentation, publicly available data are exceedingly scarce, leading to relatively small sample sizes for these six datasets.

**Fig. 2.**
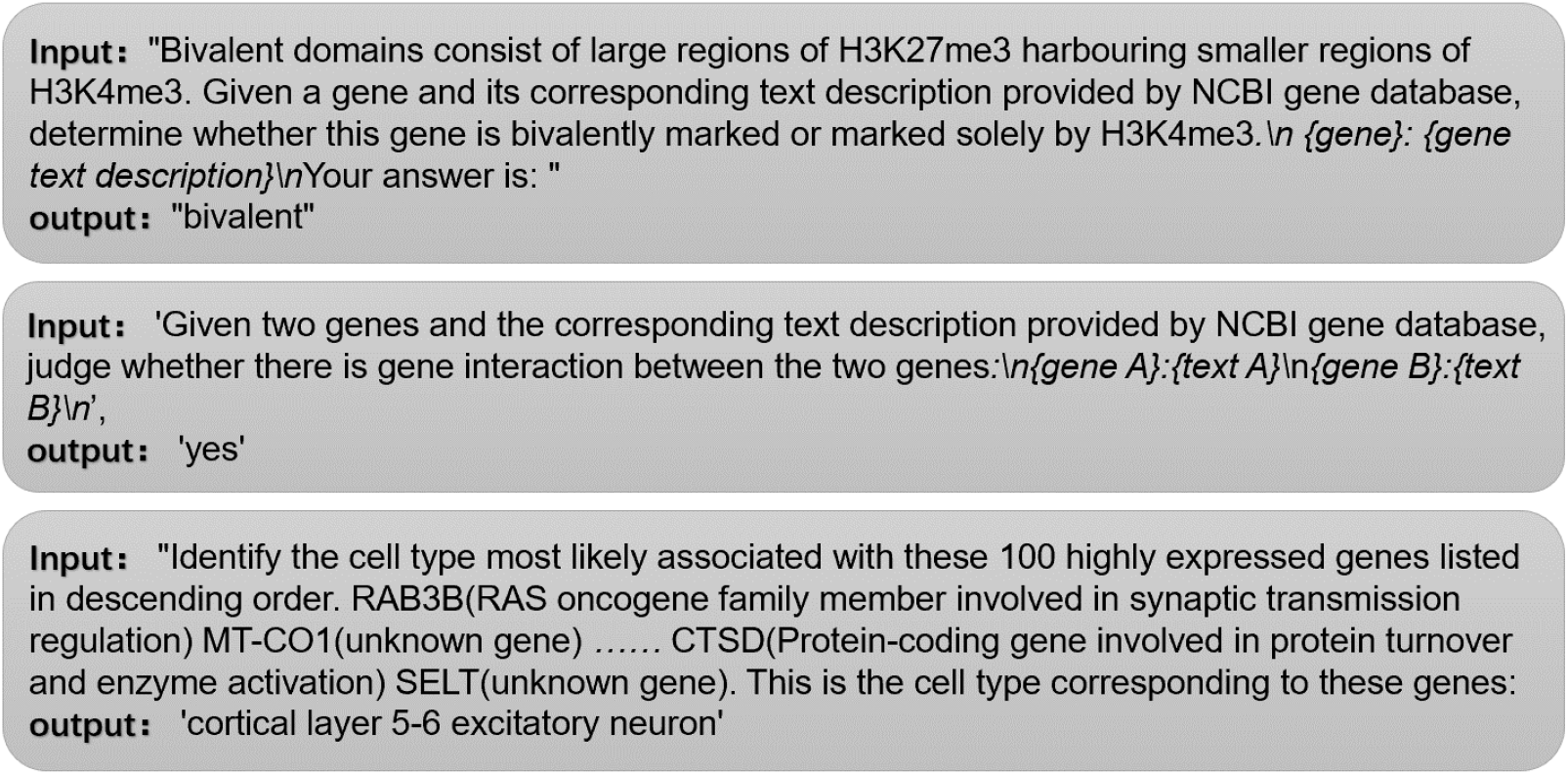
Examples of prompts for fine-tuning LLMs on three types of tasks. From top to bottom, they are: Identification of Genes, Gene-Gene Interaction, Cell Annotation.

##### Gene dosage sensitivity predictions

The dosage sensitivity of a gene refers to individuals’ sensitivity to changes in gene dosage, such as an increase in gene copy number. Some genes, when their copy number changes, can lead to significant health or developmental issues, while others may be relatively insensitive to such changes. Studying gene dosage sensitivity is crucial for genetic diagnosis. We finetuned LLMs using a dataset of 481 samples reported in prior studies[21, 29, 49] to distinguish dosage-sensitive genes and insensitive ones, divided into training and test sets at a 9:1 ratio.

##### Bivalent chromatin structure predictions

The bivalent structure of genes refers to a special chromatin structure observed in certain cell types, especially embryonic stem cells, characterized by the simultaneous presence of markers indicative of activation and repression, signifying a gene’s state of being both activated and repressed. Investigating gene bivalent structures is significant for understanding cellular characteristics and differentiation mechanisms. Our data derive from genes found in 56 conserved regions of the genome[2]. Initially, we explored whether LLMs could differentiate between bivalent genes and non-methylated genes, then further distinguished between bivalent genes and Lys4-only modified genes (indicating an activated state). The dataset sizes were 147 and 184, respectively, both divided at a 9:1 ratio.

##### Genomic distances of Transcription Factors

Transcription factors(TFs) exert their effects by binding to DNA sequences near gene promoters. TFs binding close to their target sites generally represent a rapid and direct regulatory mechanism, while long-range TFs involve more complex regulatory schemes. We utilized a dataset from Chen et al.[4], comprising 173 samples divided at a 9:1 ratio.

##### Core regulatory elements predictions

The gene regulatory mechanism is a complex process by which organisms control the timing and extent of gene expression. Identifying core regulatory elements can help pinpoint the root causes of diseases. We employed data from an NOTCH1 (N1)-dependent gene network provided by Theodoris, C. V. et al.[31, 33] to determine which genes are core elements of this N1 network and which are peripheral downstream effectors. The dataset contained 281 samples.

##### N1 downstream targets predictions

Utilizing a dataset also provided by [31, 33], we further finetuned LLMs to predict genes directly affected by this N1 network. The dataset for N1 downstream targets versus non-targets comprised 1103 samples.

#### 3.2.2 Gene Gene Interaction

The second category of tasks explores whether LLMs can predict interactions between genes. We supplied LLMs with pairs of genes to test, along with their respective NCBI records, and fine-tuned the LLMs to output predictions, as shown in Figure 2. We used the Gene-Gene Interaction(GGI) dataset from Du et al.[7], generated based on gene ontology annotations. This dataset had its training and test sets predefined, with the training set containing 249,281 samples and the test set comprising 20,338 samples after our filtration step.

#### 3.2.3 Cell Annotation

Cells serve as containers for genes, providing the platform on which genes play their role. The selective expression of genes determines the morphology and function of cells. The third category of tasks focuses on cell-level application scenarios. We selected a classic problem in the single-cell domain for evaluation: cell type annotation, a crucial step in single-cell data analysis essential for understanding the functions and characteristics of different cells. We use the Multiple Sclerosis (M.S.) dataset for experiments, initially published by Schirmer et al.[28], sampled from 9 healthy individuals and 12 patients. Following the split method of scGPT[6], we used samples from healthy individuals as the training set, totaling 7,844 samples, and samples from patients as the test set, totaling 13,316 samples. This split method introduced noise between the training and test sets, posing a challenge for the annotation models. In addition to public datasets, we also tested on data produced in our wet lab experiments, derived from human early embryos, comprising 100,321 samples across 61 cell types, divided into training and test sets at a 9:1 ratio.

Above we have introduced the basic situation of all the test tasks selected in this paper and the reasons for choosing. Table 1 summarizes the datasets used in all experiments, and Figure 2 provides an example prompt for each of the three categories of tasks.

**Table 1.** Overview of Downstream Tasks and their Datasets.

| dataset | task type | categories | train size | test size |
| --- | --- | --- | --- | --- |
| dosage sensitivity | Gene-level | 2 | 432 | 49 |
| bivalent vs non-methylated | Gene-level | 2 | 132 | 15 |
| bivalent vs lys-4 methylated | Gene-level | 2 | 165 | 19 |
| long- vs short- range TFs | Gene-level | 2 | 155 | 18 |
| N1 central vs peripheral | Gene-level | 2 | 252 | 29 |
| N1 activated vs non-target | Gene-level | 2 | 992 | 111 |
| gene gene interaction | Gene pairs level | 2 | 249281 | 20338 |
| Multiple Sclerosis cellAnno | Cell-level | 18 | 7844 | 13316 |
| hEmbyro cellAnno | Cell-level | 61 | 90288 | 10033 |

## 4 RESULTS

In this study, we selected the currently most popular LLMs as the subjects for our evaluation, including Llama, Mistral, BLOOM, and ChatGLM. The rationale behind this selection is discussed in the first paragraph of Chapter 2. For each of these LLMs, the 7-billion-parameter version was chosen for evaluation, with Llama-13b serving as the representative for larger parameter models. Given that ChatGPT’s latest model is not open-sourced, we included its predecessor, GPT-2, in our evaluation. BioGPT, a specialized LLM for the biomedical literature domain, was also included in this evaluation. All LLMs were uniformly fine-tuned using LORA, with details on the parameter count and pre-training data volume presented in Table 2.

**Table 2.** Overview of the LLMs to be evaluated and their final score.

| LLMS | trainable params | all params | pre-train dataset size | final scores | rank |
| --- | --- | --- | --- | --- | --- |
| GPT2 | 294,912 | 124,734,720 | 40G | 0.568 | 7 |
| BioGPT | 786,432 | 347,549,696 | 15M | 0.721 | 6 |
| ChatGLM | 1,949,696 | 6,245,533,696 | 1T | 0.741 | 4 |
| BLOOM | 3,932,160 | 7,072,948,224 | 1.61T | 0.760 | 2 |
| Mistral | 3,407,872 | 7,245,139,968 | 8T | 0.754 | 3 |
| Llama-7b | 4,194,304 | 6,742,609,920 | 1.4T | 0.736 | 5 |
| Llama-13b | 6,553,600 | 13,022,417,920 | 1.4T | 0.768 | 1 |

### 4.1 Through fine-tuning, LLMs can understand genes from different perspectives

Recognizing that datasets related to the Identification of genes functions and traits are generally small in size, as shown in Table 1, we demonstrated the LLMs’ test results using the number of correctly predicted test cases and calculated the overall accuracy across six datasets as the score for this task category. The final results, as shown in Table 3, are presented succinctly, with dataset names abbreviated and corresponding to those listed in Table 1 in sequence. It was observed that Llama-13b performed the best, affirming that model parameter size remains a crucial factor in gene-related domains. BLOOM and BioGPT followed closely behind, ranking second and third respectively, suggesting that the diversity (BLOOM was pre-trained on multilingual corpora) and specialization (BioGPT was pre-trained on specialized biological corpora) of pre-training corpora significantly contribute to model performance. These conclusions are in line with experiences from other scientific fields where LLMs have been applied.

**Table 3.** Results of tasks related to Identification of Genes, measured by the correct number of predictions.

| LLMS | dosage | methyalted | lys-4 | range | N1 central | N1 activated | accuracy |
| --- | --- | --- | --- | --- | --- | --- | --- |
| GPT2 | 39 | 11 | 15 | 13 | 19 | 68 | 0.684 |
| BioGPT | 41 | 14 | 16 | 10 | 25 | **80** | 0.772 |
| ChatGLM | 40 | 13 | 17 | 13 | 20 | 78 | 0.751 |
| BLOOM | 41 | 14 | **18** | 12 | 23 | **80** | 0.780 |
| Mistral | **43** | **15** | **18** | 13 | 21 | 73 | 0.759 |
| Llama-7b | 40 | 12 | 17 | 13 | 22 | 70 | 0.722 |
| Llama-13b | **43** | 13 | 17 | 12 | **26** | 78 | **0.784** |

Moreover, in experiments to identify whether genes are bivalent or unmethylated, Mistral achieved the best results, correctly predicting all 15 test cases. Notably, only Mistral successfully predicted a special case of TCF12 among all models. TCF12 is a transcription factor, and in our dataset, most TFs were bivalently modified (out of 15 test cases, 12 were TFs, with only TCF12 being unmethylated). In addition, Mistral has a total of five cases in which only its own prediction is correct and other models are wrong, with the highest number of all LLMs, followed by BioGPT (three cases) and Llama-13b(two cases). Mistral’s exceptional performance on more challenging test cases can likely be attributed to its extensive pre-training on 8 trillion tokens, allowing it to be exposed to a broader range of corpora than the other models.

In experiments distinguishing long-range and short-range TFs, all models incorrectly predicted the SPIB gene. This may be attributed to the descriptions provided to the LLMs, which mentioned that the SPIB gene is a lymphocyte-specific enhancer, suggesting a regulatory relationship requiring rapid response to external signals, thus leading all models to classify it as a short-range transcription factor. However, Tfs often do not have a singular mode of action, and SPIB could potentially interact with other proteins through long-range mechanisms under certain conditions, as indicated by its experimental determination as long-range. Such scenarios are not uncommon in the biological literature. Since much biological knowledge is derived from experimental findings, which may involve randomness or errors, the biological corpora could contain biased or incorrect descriptions, potentially leading to partial or erroneous outputs from LLMs in certain applications.

### 4.2 LLMs is able to predict most gene pairs interrelation

The performance of various LLMs on the GGI task is presented in Table 4. GPT-2’s performance was notably inferior to that of other models, likely due to its smaller number of trainable parameters which may have limited its ability to fit such extensive data. The results for Mistral, BLOOM, Llama-7b, and Llama-13b followed a similar pattern where a larger number of trainable parameters correlated with improved performance. Exceptions to this trend were ChatGLM and BioGPT. ChatGLM, with less than two million parameters, outperformed Llama-7b, which possesses over four million parameters, earning it notable praise. BioGPT, leveraging its outdated model architecture with less than one million parameters, surpassed all other models, exceeding the next best model (Llama-13b) by 6.5 percentage points. The success of BioGPT on the GGI task underscores the importance of domain-specific pre-training in the field of single-cell research.

**Table 4.**
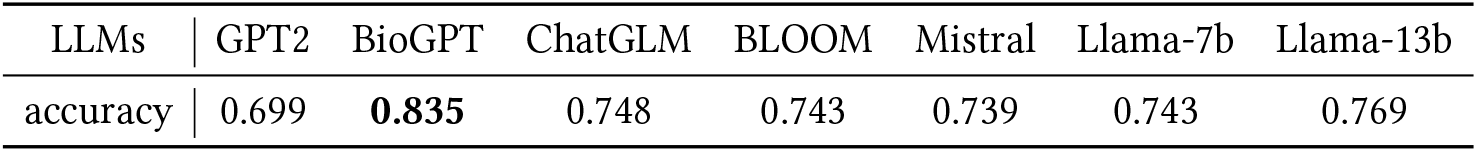
Results of GGI task, measured by prediction accuracy.

| LLMs | GPT2 | BioGPT | ChatGLM | BLOOM | Mistral | Llama-7b | Llama-13b |
| --- | --- | --- | --- | --- | --- | --- | --- |
| accuracy | 0.699 | **0.835** | 0.748 | 0.743 | 0.739 | 0.743 | 0.769 |

Within the GGI test set, encompassing 20,338 instances, nearly half (9,841 instances) were accurately predicted by all models, with only a small fraction(734 instances) incorrectly predicted across the board. This outcome is reasonable, because the GGI dataset is derived from genes’ commonalities in their ontology annotations. Gene Ontology(GO) is a system for describing genes using a set of terms, akin to tagging genes with appropriate labels based on their functions. The gene’s NCBI records and its GO annotation offer complementary perspectives, yet for many genes, these descriptions are substantially overlapping (even some genes contain their GO annotation terms in their NCBI records). For such genes, LLMs is easy to judge whether they have commonality according to NCBI records, whereas the prediction of other less straightforward cases challenges the models’ capacity for knowledge extrapolation. Among all the LLMs evaluated, BioGPT achieved the best and far leading performance on the GGI task, a feat not replicated in other downstream tasks explored within this paper. This can likely be attributed to its pre-training on biological literature, which would have exposed it to a large corpus of NCBI descriptions and GO annotations.

### 4.2 LLMs’ Ability in Cell Annotation Tasks

The performance of all LLMs on cell annotation tasks is shown in Table 5. The left side presents results for the M.S. dataset (comprising 18 categories), and the right side for the human embryonic dataset (comprising 61 categories). Given that the M.S. dataset’s train and test sets originate from different individuals, predicting cell types presents a higher challenge. We employed traditional multi-classification metrics to assess model performance. Notably, the accuracy metric is calculated globally (macro), and the remaining three metrics are calculated by categories average (micro). The results demonstrate that Mistral achieved the best performance on both datasets, closely followed by BLOOM. Considering the annotation task involved processing long token sequences (averaging about 2,000 tokens), Mistral 8T’s extensive pre-training data volume enhanced its understanding of long texts. Conversely, GPT-2 and BioGPT showed significant performance drops, primarily due to their smaller model and pre-training dataset sizes, limiting their capability with long text inputs. To validate the effectiveness of the “cell sentence plus” approach, we also conducted an ablation study by traditional “cell sentence” methods to textually represent cells, followed by training large models using the same prompts. As shown in Table 6, the “cell sentence plus” technique improved performance for all LLMs above 6 billion parameters, affirming the validity of our approach. However, for GPT-2 and BioGPT, lacking the capability to process long texts, the “cell sentence plus” method proved to be burdensome.

**Table 5.** Results of Cell Annotaion tasks, measured by traditional classification metrics.

|  | Multiple Sclerosis |  |  |  | hEmbryo |  |  |  |
| --- | --- | --- | --- | --- | --- | --- | --- | --- |
| LLMS | accuracy | recall | precision | F1 score | accuracy | recall | precision | F1 score |
| GPT2 | 0.355 | 0.271 | 0.289 | 0.251 | 0.288 | 0.191 | 0.224 | 0.189 |
| BioGPT | 0.623 | 0.491 | 0.539 | 0.471 | 0.586 | 0.500 | 0.550 | 0.510 |
| ChatGLM | 0.678 | 0.520 | 0.569 | 0.507 | 0.771 | 0.732 | 0.768 | 0.747 |
| BLOOM | 0.714 | 0.571 | **0.605** | 0.557 | 0.797 | 0.769 | **0.794** | 0.78 |
| Mistral | **0.715** | **0.586** | 0.595 | **0.577** | **0.814** | 0.775 | 0.792 | **0.782** |
| Llama-7b | 0.686 | 0.553 | 0.587 | 0.546 | 0.799 | **0.776** | 0.792 | **0.782** |
| Llama-13b | 0.701 | 0.582 | 0.593 | 0.569 | 0.802 | 0.773 | 0.789 | 0.780 |

**Table 6.** Ablation Study: Cell Sentence vs Cell Sentence Plus.

| Model | Multiple Sclerosis |  |  |  | hEmbyro |  |  |  |
| --- | --- | --- | --- | --- | --- | --- | --- | --- |
|  | acc | acc(plus) ↑↓ | F1 | F1(plus) ↑↓ | acc | acc(plus) ↑↓ | F1 | F1(plus) ↑↓ |
| GPT2 | 0.526 | 0.355 (↓0.171) | 0.377 | 0.251 (↓0.126) | 0.266 | 0.288 (↑0.022) | 0.156 | 0.189 (↑0.033) |
| BioGPT | 0.640 | 0.623 (↓0.017) | 0.480 | 0.471 (↓0.009) | 0.590 | 0.586 (↓0.004) | 0.524 | 0.510 (↓0.014) |
| ChatGLM | 0.644 | 0.678 (↑0.034) | 0.495 | 0.507 (↑0.012) | 0.743 | 0.771 (↑0.028) | 0.715 | 0.747 (↑0.032) |
| BLOOM | 0.693 | 0.714 (↑0.021) | 0.536 | 0.557 (↑0.021) | 0.774 | 0.797 (↑0.023) | 0.749 | 0.780 (↑0.031) |
| Mistral | 0.707 | **0.715** (↑0.008) | 0.569 | **0.577** (↑0.008) | 0.807 | **0.814** (↑0.007) | 0.776 | **0.782** (↑0.006) |
| Llama-7b | 0.685 | 0.686 (↑0.001) | 0.526 | 0.546 (↑0.020) | 0.779 | 0.799 (↑0.020) | 0.757 | **0.782** (↑0.025) |
| Llama-13b | 0.688 | 0.701 (↑0.013) | 0.540 | 0.569 (↑0.029) | 0.789 | 0.802 (↑0.013) | 0.761 | 0.780 (↑0.019) |

We took the average of each model’s prediction accuracy over the two datasets as their score on this type of task. The average scores of the three types of tasks are calculated as the final score of LLMs evaluation. The final scores and their rankings are shown in Table 2, with the only 13b LLM taking the first place, with BLOOM and Mistral in second and third place. Due to its smaller parameter size, BioGPT ranked lower in cell-level tasks, yet its performance in the first two task categories was exceptional, underscoring the impact of domain-specific large models in solving niche area problems. We look forward to the emergence of biological domain LLMs built on the latest model architectures.

## 5 DISCUSSION

In this paper, we demonstrate how fine-tuning LLMs can address issues related to genes and cells, and we evaluate the performance of seven LLMs across nine datasets. The scenarios encompass tasks such as identifying individual gene identities, predicting relationships between gene pairs, and cell-level annotation. The 13b version of Llama, as the sole 13-billion-parameter model in our study, emerged as the frontrunner, underscoring the significance of model size for the efficacy of LLMs. BLOOM and Mistral, ranking second and third, highlight the enhancement in LLM performance attributed to the diversity and volume of pre-training corpora. BioGPT, despite its smaller scale and inability to process long text inputs compared to other models, showcased a remarkable understanding of genes and GGI relationships, pointing to the importance of domain-specific pre-training corpora. Overall, LLMs performed commendably across all tasks, illustrating their potential to comprehend and even solve gene-related issues to a certain extent. However, in specific tasks like cell type annotation, LLMs’ predictions still fall short when compared to specialized single-cell sequencing models. On the M.S. dataset, LLMs achieved an accuracy rate of about 0.7, whereas specialized models could reach above 0.8. Nevertheless, considering the limited input information (only the top 100 gene names) and lower training costs for LLMs, such outcomes are acceptable. Currently, specialized models either involve complex modeling of gene relationships or require pre-training on large-scale sequencing data, whereas fine-tuning LLMs merely necessitates considering to conceptualize the issue in textual form.

Applying LLMs in cellular biology presents several challenges. Firstly, the scarcity of biomedically relevant textual corpora limits the development of domain-specific large parameter LLMs. Secondly, the presence of bias and even erroneous descriptions in biological literature could potentially mislead LLMs into making incorrect judgments. Furthermore, biological processes often involve complex regulatory and causal relationships, with many mechanisms still beyond our current understanding. Much of this information is difficult to convey through text alone, restricting LLMs to solving only relatively superficial biological problems. However, given their flexible interaction modes and powerful capabilities in information retrieval and synthesis, LLMs can assist cellular biologists in quickly acquiring basic knowledge about genes and cells and provide preliminary insights into more complex issues. We recommend researchers in gene-related fields use LLMs as tools for information retrieval and idea generation, rather than relying solely on LLM conclusions to address practical problems.

Our work also has its limitations. Our evaluation scope is not comprehensive enough, not only for the LLMs but also for selected downstream tasks. Our evaluation metrics are too simplistic, considering only the accuracy of fine-tuned LLMs in solving gene-related classification tasks and neglecting other aspects such as training efficiency, hyperparameter selection, and robustness, which only crudely reflect the potential application of LLMs in this field. In the future, we will attempt to conduct evaluations using more professional and comprehensive methods[8, 17, 37, 47]. Our prompts for LLMs are also quite straightforward, requiring a more thoughtful design for practical problem-solving.

Regarding the future direction of applying LLMs to gene-related issues, we offer the following suggestions: 1) Construct specialized LLMs for the biological domain based on the latest LLM architectures. The main challenge lies in the scarcity of biological corpora. Apart from collecting more corpora, approaches such as data augmentation and adversarial learning should be considered. 2) Develop cell sentence pre-trained LLMs. High-throughput sequencing technology has accumulated billions of transcriptomic data points, sufficient for large model pre-training. Processing these data into a cell sentence plus format for LLM pre-training could enhance the understanding of information encapsulated in this format, aiding in addressing many downstream tasks related to sequencing data. 3) Explore more downstream scenarios for LLMs. The tasks involved in this paper tend to focus on the understanding of genes and cells. For more specific application scenarios, such as embryonic development, gene editing, and the genetics of aging, LLMs are also expected to play a larger role, but how to textually describe these problems is a challenge. Moreover, due to the costly nature of biological experiments, experimental data in many biological contexts is extremely limited. To alleviate this, we consider introducing semi-supervised learning[18, 24, 35, 36, 43, 48] methods to tackle gene-related problems, which aim to enhance the limited labeled data with large-scale unlabeled data. Additionally, for certain downstream tasks where data imbalance is a severe issue, we consider introducing methods from Imbalanced Learning[38–40, 44, 45] to increase focus on rare categories. All in all, LLMs can not only serve as a research tool for cell biology researchers, but also is expected to play a greater role in solving gene-related problems.

## REFERENCES

[1] T. Barrett, S. E. Wilhite, P. Ledoux, C. Evangelista, I. F. Kim, M. Tomashevsky, K. A. Marshall, K. H. Phillippy, P. M. Sherman, M. Holko, A. Yefanov, H. Lee, N. Zhang, C. L. Robertson, N. Serova, S. Davis, and A. Soboleva. 2013. NCBI GEO: archive for functional genomics data sets–update. Nucleic Acids Res 41, Database issue (Jan 2013), D991–995.

[2] Bradley E. Bernstein, Tarjei S. Mikkelsen, Xiaohui Xie, Michael Kamal, Dana J. Huebert, James Cuff, Ben Fry, Alex Meissner, Marius Wernig, Kathrin Plath, Rudolf Jaenisch, Alexandre Wagschal, Robert Feil, Stuart L. Schreiber, and Eric S. Lander. 2006. A Bivalent Chromatin Structure Marks Key Developmental Genes in Embryonic Stem Cells. Cell 125, 2 (2006), 315–326. 10.1016/j.cell.2006.02.041

[3] Garth R. Brown, Vichet Hem, Kenneth S. Katz, Michael Ovetsky, Craig Wallin, Olga Ermolaeva, Igor Tolstoy, Tatiana Tatusova, Kim D. Pruitt, Donna R. Maglott, and Terence D. Murphy. 2014. Gene: a gene-centered information resource at NCBI. Nucleic Acids Research 43, D1 (10 2014), D36–D42. 10.1093/nar/gku1055 arXiv:https://academic.oup.com/nar/article-pdf/43/D1/D36/7314925/gku1055.pdf

[4] Chenhao Chen, Rongbin Zheng, Collin J Tokheim, Xin Dong, Jingyu Fan, Changxin Wan, Qin Tang, Myles A. Brown, Jun S. Liu, Clifford A. Meyer, and Shirley X. Liu. 2019. Determinants of transcription factor regulatory range. Nature Communications 11 (2019). https://api.semanticscholar.org/CorpusID:91596152

[5] Y. T. Chen and J. Zou. 2023. GenePT: A Simple But Hard-to-Beat Foundation Model for Genes and Cells Built From ChatGPT. bioRxiv (Oct 2023).

[6] Haotian Cui, Chloe Wang, Hassaan Maan, and Bo Wang. [n. d.]. scGPT: Towards Building a Foundation Model for Single-Cell Multi-omics Using Generative AI. ([n. d.]). 10.1101/2023.04.30.538439

[7] Jingcheng Du, Peilin Jia, Yulin Dai, Cui Tao, Zhongming Zhao, and Degui Zhi. 2019. Gene2vec: Distributed representation of genes based on co-expression. BMC Genomics 20 (02 2019). 10.1186/s12864-018-5370-x

[8] Leo Gao, Jonathan Tow, Baber Abbasi, Stella Biderman, Sid Black, Anthony DiPofi, Charles Foster, Laurence Golding, Jeffrey Hsu, Alain L. Noac’h, Haonan Li, Kyle McDonell, Niklas Muennighoff, Chris Ociepa, Jason Phang, Laria Reynolds, Hailey Schoelkopf, Aviya Skowron, Lintang Sutawika, Eric Tang, Anish Thite, Ben Wang, Kevin Wang, and Andy Zou. 2023. A framework for few-shot language model evaluation. 10.5281/zenodo.10256836

[9] Minsheng Hao, Jing Gong, Xin Zeng, Chiming Liu, Yucheng Guo, Xingyi Cheng, Taifeng Wang, Jianzhu Ma, L. Song, and Xuegong Zhang. 2023. Large Scale Foundation Model on Single-cell Transcriptomics. bioRxiv (2023). 10.1101/2023.05.29.542705 arXiv:https://www.biorxiv.org/content/early/2023/05/31/2023.05.29.542705.full.pdf

[10] hiyouga. 2023. LLaMA Factory. https://github.com/hiyouga/LLaMA-Factory.

[11] W. Hou and Z. Ji. 2023. Assessing GPT-4 for cell type annotation in single-cell RNA-seq analysis. bioRxiv (Dec 2023).

[12] Edward J. Hu, Yelong Shen, Phillip Wallis, Zeyuan Allen-Zhu, Yuanzhi Li, Shean Wang, Lu Wang, and Weizhu Chen. 2021. LoRA: Low-Rank Adaptation of Large Language Models. arXiv:2106.09685 [cs.CL]

[13] Byungjin Hwang, Ji Hyun Lee, and Duhee Bang. 2018. Single-cell RNA sequencing technologies and bioinformatics pipelines. Experimental & Molecular Medicine 50 (2018). https://api.semanticscholar.org/CorpusID:51942532

[14] Albert Q. Jiang, Alexandre Sablayrolles, Arthur Mensch, Chris Bamford, Devendra Singh Chaplot, Diego de las Casas, Florian Bressand, Gianna Lengyel, Guillaume Lample, Lucile Saulnier, Lélio Renard Lavaud, Marie-Anne Lachaux, Pierre Stock, Teven Le Scao, Thibaut Lavril, Thomas Wang, Timothée Lacroix, and William El Sayed. 2023. Mistral 7B. arXiv:2310.06825 [cs.CL]

[15] Dragomirka Jovic, Xue Liang, Hua Zeng, Lin Lin, Fengping Xu, and Yonglun Luo. 2022. Single-cell RNA sequencing technologies and applications: A brief overview. Clinical and Translational Medicine 12 (03 2022). 10.1002/ctm2.694

[16] Daniel Levine, Sacha Lévy, Syed Asad Rizvi, Nazreen Pallikkavaliyaveetil, Xingyu Chen, et al. 2024. Cell2Sentence: Teaching Large Language Models the Language of Biology. bioRxiv (2024). 10.1101/2023.09.11.557287 arXiv:https://www.biorxiv.org/content/early/2024/02/15/2023.09.11.557287.full.pdf

[17] Qingqing Long, Yilun Jin, Yi Wu, and Guojie Song. 2021. Theoretically improving graph neural networks via anonymous walk graph kernels. In Proceedings of the Web Conference 2021. 1204–1214.

[18] Qingqing Long, Lingjun Xu, Zheng Fang, and Guojie Song. 2021. Hgk-gnn: heterogeneous graph kernel based graph neural networks. In Proceedings of the 27th ACM SIGKDD Conference on Knowledge Discovery & Data Mining. 1129–1138.

[19] Romain Lopez, Jeffrey Regier, Michael B. Cole, Michael I. Jordan, and Nir Yosef. [n. d.]. Deep generative modeling for single-cell transcriptomics. 15, 12 ([n. d.]), 1053–1058. 10.1038/s41592-018-0229-2

[20] Renqian Luo, Liai Sun, Yingce Xia, Tao Qin, Sheng Zhang, Hoifung Poon, and Tie-Yan Liu. 2022. BioGPT: generative pre-trained transformer for biomedical text generation and mining. Briefings in Bioinformatics 23, 6 (09 2022), bbac409. 10.1093/bib/bbac409 arXiv:https://academic.oup.com/bib/article-pdf/23/6/bbac409/47144271/bbac409.pdf

[21] Zhihua Ni, Xiao-Yu Zhou, Sidra Aslam, and Deng-Ke Niu. 2019. Characterization of Human Dosage-Sensitive Transcription Factor Genes. Frontiers in Genetics 10 (12 2019), 1208. 10.3389/fgene.2019.01208

[22] OpenAI. 2023. GPT-4 Technical Report. arXiv:2303.08774 [cs.CL]

[23] Alec Radford and Karthik Narasimhan. 2018. Improving Language Understanding by Generative Pre-Training. https://api.semanticscholar.org/CorpusID:49313245

[24] Y. C. A. Padmanabha Reddy, P. Viswanath, and B. Eswara Reddy. 2018. Semi-supervised learning: a brief review. International journal of engineering and technology 7 (2018), 81. https://api.semanticscholar.org/CorpusID:55044284

[25] A. Regev, S. A. Teichmann, E. S. Lander, I. Amit, C. Benoist, E. Birney, B. Bodenmiller, P. Campbell, et al. 2017. The Human Cell Atlas. Elife 6 (Dec 2017).

[26] Yusuf Roohani, Kexin Huang, and Jure Leskovec. 2023. Predicting transcriptional outcomes of novel multigene perturbations with GEARS. Nature Biotechnology (17 Aug 2023). 10.1038/s41587-023-01905-6

[27] Teven Le Scao, Angela Fan, Christopher Akiki, Ellie Pavlick, Suzana Ili’c, Daniel Hesslow, Roman Castagn’e, et al. 2022. BLOOM: A 176B-Parameter Open-Access Multilingual Language Model. ArXiv abs/2211.05100 (2022). https://api.semanticscholar.org/CorpusID:253420279

[28] Lucas Schirmer, Dmitry Velmeshev, Staffan Holmqvist, Max Kaufmann, Sebastian Werneburg, Diane Jung, Stephanie Vistnes, John Stockley, Adam Young, Maike Steindel, Brian Tung, Nitasha Goyal, Aparna Bhaduri, Simone Mayer, Jan Engler, Omer Ali Bayraktar, Robin Franklin, Maximilian Haeussler, Richard Reynolds, and David Rowitch. 2019. Neuronal vulnerability and multilineage diversity in multiple sclerosis. Nature 573 (09 2019), 1. 10.1038/s41586-019-1404-z

[29] Hashem A Shihab, Mark F Rogers, Colin Campbell, and Tom R Gaunt. 2017. HIPred: an integrative approach to predicting haploinsufficient genes. Bioinformatics 33, (01 2017), 1751–1757. 10.1093/bioinformatics/btx028 arXiv:https://academic.oup.com/bioinformatics/article-pdf/33/12/1751/49039929/bioinformatics_33_12_1751.pdf

[30] Hantao Shu, Jingtian Zhou, Qiuyu Lian, Han Li, Dan Zhao, Jianyang Zeng, and Jianzhu Ma. 2021. Modeling gene regulatory networks using neural network architectures. Nature Computational Science 1, 7 (01 Jul 2021), 491–501. 10.1038/s43588-021-00099-8

[31] Christina V. Theodoris, Molong Li, Mark P. White, Lei Liu, Daniel He, Katherine S. Pollard, Benoit G. Bruneau, and Deepak Srivastava. 2015. Human Disease Modeling Reveals Integrated Transcriptional and Epigenetic Mechanisms of NOTCH1 Haploinsufficiency. Cell 160, 6 (2015), 1072–1086. 10.1016/j.cell.2015.02.035

[32] Christina V. Theodoris, Ling Xiao, Anant Chopra, Mark D. Chaffin, Zeina R. Al Sayed, Matthew C. Hill, Helene Mantineo, Elizabeth M. Brydon, Zexian Zeng, X. Shirley Liu, and Patrick T. Ellinor. [n. d.]. Transfer learning enables predictions in network biology. 618, 7965 ([n. d.]), 616–624. 10.1038/s41586-023-06139-9

[33] Christina V. Theodoris, Ping Zhou, Lei Liu, Yu Zhang, Tomohiro Nishino, Yu Huang, Aleksandra Kostina, Sanjeev S. Ranade, Casey A. Gifford, Vladimir Uspenskiy, Anna Malashicheva, Sheng Ding, and Deepak Srivastava. 2021. Network-based screen in iPSC-derived cells reveals therapeutic candidate for heart valve disease. Science 371, 6530 (2021), eabd0724. 10.1126/science.abd0724 arXiv:https://www.science.org/doi/pdf/10.1126/science.abd0724

[34] Hugo Touvron, Louis Martin, Kevin Stone, Peter Albert, Amjad Almahairi, Yasmine Babaei, Nikolay Bashlykov, et al. 2023. Llama 2: Open Foundation and Fine-Tuned Chat Models. arXiv:2307.09288 [cs.CL]

[35] Yidong Wang, Hao Chen, Yue Fan, Wang SUN, Ran Tao, Wenxin Hou, Renjie Wang, Linyi Yang, Zhi Zhou, Lan-Zhe Guo, Heli Qi, Zhen Wu, Yu-Feng Li, Satoshi Nakamura, Wei Ye, Marios Savvides, Bhiksha Raj, Takahiro Shinozaki, Bernt Schiele, Jindong Wang, Xing Xie, and Yue Zhang. 2022. USB: A Unified Semi-supervised Learning Benchmark for Classification.In Advances in Neural Information Processing Systems, S. Koyejo, S. Mohamed, A. Agarwal, D. Belgrave, K. Cho, and A. Oh (Eds.), Vol. 35. Curran Associates, Inc., 3938–3961. https://proceedings.neurips.cc/paper_files/paper/ 2022/file/190dd6a5735822f05646dc27decff19b-Paper-Datasets_and_Benchmarks.pdf

[36] Yidong Wang, Hao Chen, Qiang Heng, Wenxin Hou, Yue Fan,, Zhen Wu, Jindong Wang, Marios Savvides, Takahiro Shinozaki, Bhiksha Raj, Bernt Schiele, and Xing Xie. 2023. FreeMatch: Self-adaptive Thresholding for Semi-supervised Learning. (2023).

[37] Yidong Wang, Zhuohao Yu, Zhengran Zeng, Linyi Yang, Cunxiang Wang, Hao Chen, Chaoya Jiang, Rui Xie, Jindong Wang, Xing Xie, Wei Ye, Shikun Zhang, and Yue Zhang. 2024. PandaLM: An Automatic Evaluation Benchmark for LLM Instruction Tuning Optimization. (2024).

[38] Yidong Wang, Bowen Zhang, Wenxin Hou, Zhen Wu, Jindong Wang, and Takahiro Shinozaki. 2023. Margin Calibration for Long-Tailed Visual Recognition. In Proceedings of The 14th Asian Conference on Machine Learning (Proceedings of Machine Learning Research, Vol. 189), Emtiyaz Khan and Mehmet Gonen (Eds.). PMLR, 1101–1116. https://proceedings.mlr.press/v189/wang23b.html

[39] Yidong Wang, Yu Zhuohao, Jindong Wang, Qiang Heng, Hao Chen, Wei Ye, Xie Rui, Xing Xie, and Shikun Zhang. 2023. Exploring Vision-Language Models for Imbalanced Learning. International Journal of Computer Vision 132 (08 2023). 10.1007/s11263-023-01868-w

[40] Yu-Xiong Wang, Deva Ramanan, and Martial Hebert. 2017. Learning to model the tail (NIPS’17). Curran Associates Inc., Red Hook, NY, USA, 7032–7042.

[41] Yuchen Yan, Peiyan Zhang, Zheng Fang, and Qingqing Long. 2024. Inductive Graph Alignment Prompt: Bridging the Gap between Graph Pre-training and Inductive Fine-tuning From Spectral Perspective. arXiv preprint arXiv:2402.13556 (2024).

[42] Xiaodong Yang, Guole Liu, Guihai Feng, Dechao Bu, Pengfei Wang, Jie Jiang, Shubai Chen, et al. 2023. GeneCompass: Deciphering Universal Gene Regulatory Mechanisms with Knowledge-Informed Cross-Species Foundation Model. bioRxiv (2023). 10.1101/2023.09.26.559542 arXiv:https://www.biorxiv.org/content/early/2023/09/28/2023.09.26.559542.full.pdf

[43] Xiangli Yang, Zixing Song, Irwin King, and Zenglin Xu. 2023. A Survey on Deep Semi-Supervised Learning. IEEE Transactions on Knowledge and Data Engineering 35, 9 (2023), 8934–8954. 10.1109/TKDE.2022.3220219

[44] Yuzhe Yang and Zhi Xu. 2020. Rethinking the Value of Labels for Improving Class-Imbalanced Learning. arXiv:2006.07529 [cs.LG]

[45] Haiyang Yu, Ningyu Zhang, Shumin Deng, Zonggang Yuan, Yantao Jia, and Huajun Chen. 2020. The Devil is the Classifier: Investigating Long Tail Relation Classification with Decoupling Analysis. arXiv:2009.07022 [cs.LG]

[46] Aohan Zeng, Xiao Liu, Zhengxiao Du, Zihan Wang, Hanyu Lai, Ming Ding, Zhuoyi Yang, Yifan Xu, Wendi Zheng, Xiao Xia, Weng Lam Tam, Zixuan Ma, Yufei Xue, Jidong Zhai, Wenguang Chen, Peng Zhang, Yuxiao Dong, and Jie Tang. 2023. GLM-130B: An Open Bilingual Pre-trained Model. arXiv:2210.02414 [cs.CL]

[47] Lianmin Zheng, Wei-Lin Chiang, Ying Sheng, Siyuan Zhuang, Zhanghao Wu, Yonghao Zhuang, Zi Lin, Zhuohan Li, Dacheng Li, Eric. P Xing, Hao Zhang, Joseph E. Gonzalez, and Ion Stoica. 2023. Judging LLM-as-a-judge with MT-Bench and Chatbot Arena. arXiv:2306.05685 [cs.CL]

[48] Xiaojin Zhu. 2008. Semi-Supervised Learning Literature Survey. Comput Sci, University of Wisconsin-Madison 2 (07 2008).

[49] James Y. Zou. 2015. Analysis of protein-coding genetic variation in 60,706 humans. Nature 536 (2015), 285 – 291. https://api.semanticscholar.org/CorpusID:4454417

